# Performance trade-offs for single- and dual-objective light-sheet microscope designs

**DOI:** 10.1101/2020.05.10.087171

**Authors:** Kevin W. Bishop, Adam K. Glaser, Jonathan T.C. Liu

## Abstract

Light-sheet microscopy (LSM) has emerged as a powerful tool for high-speed volumetric imaging of live model organisms and large optically cleared specimens. When designing cleared-tissue LSM systems with certain desired imaging specifications (e.g. resolution, contrast, and working distance), various design parameters must be taken into consideration. In order to elucidate some of the key design trade-offs for LSM systems, we present a diffraction-based analysis of single- and dual-objective LSM configurations where Gaussian illumination is utilized. Specifically, we analyze the effects of the illumination and collection numerical aperture (NA), as well as their crossing angle, on spatial resolution and contrast. Assuming an open-top light-sheet (OTLS) architecture, we constrain these parameters based on fundamental geometric considerations as well as those imposed by currently available microscope objectives. In addition to revealing the performance tradeoffs of various single- and dual-objective LSM configurations, our analysis showcases the potential advantages of a novel, non-orthogonal dual-objective (NODO) architecture, especially for moderate-resolution imaging applications (collection NA of 0.5 to 0.8).

## 1. Introduction

Light-sheet microscopy (LSM), also known as selective plane illumination microscopy (SPIM), has become a valuable tool for many biomedical investigations and applications. In LSM, a sheet of light is used to excite fluorescence from a thin plane (“optical section”) within a relatively transparent sample [1, 2]. Adjacent 2D planes within the sample are successively illuminated and imaged using a high-speed camera to rapidly scan a volumetric region. A key advantage of this camera-based technique is that 3D imaging can be performed more quickly and simply than with laser-scanning microscopy (e.g. confocal and multiphoton microscopy). In addition, selective planar illumination is optically efficient, minimizing photobleaching of fluorophores and phototoxicity to living organisms [3]. LSM was originally popularized for volumetric imaging of live model organisms in developmental biology [1, 4–6] and more recently has been explored for imaging large optically cleared ex vivo tissues [7–9], including clinical specimens [10–12].

A number of unique LSM architectures exist that can be broadly separated into two categories: dual-objective and single-objective LSM. Dual-objective systems use two separate objectives, arranged orthogonally to one another, for illumination and collection. Having two independent objectives provides optical-design flexibility but can place significant physical constraints on sample geometries. In an attempt to mitigate this issue, inverted LSM [5, 8] and open-top light-sheet (OTLS) configurations [6,10,11,13,14] have been introduced in which dual objectives are arranged at 45° angles with respect to a sample that is placed on a horizontal substrate Fig. 1a). Such designs reduce constraints on sample size and sample numbers (by allowing for unconstrained lateral image tiling) and simplify sample mounting. However, positioning the objectives at such oblique angles typically results in a reduction in axial (vertical) imaging range since the full working distance of the objectives cannot be easily accessed (Fig. 1b). Additionally, achieving moderate- or high-numerical aperture (NA) imaging (i.e. NA > 0.5) using an OTLS architecture is challenging due to the precise index-matching requirements and/or corrective optics needed to minimize aberrations when high-NA off-axis beams transition between different media (e.g. immersion oil, sample holder, tissue) [14].

**Fig. 1.**
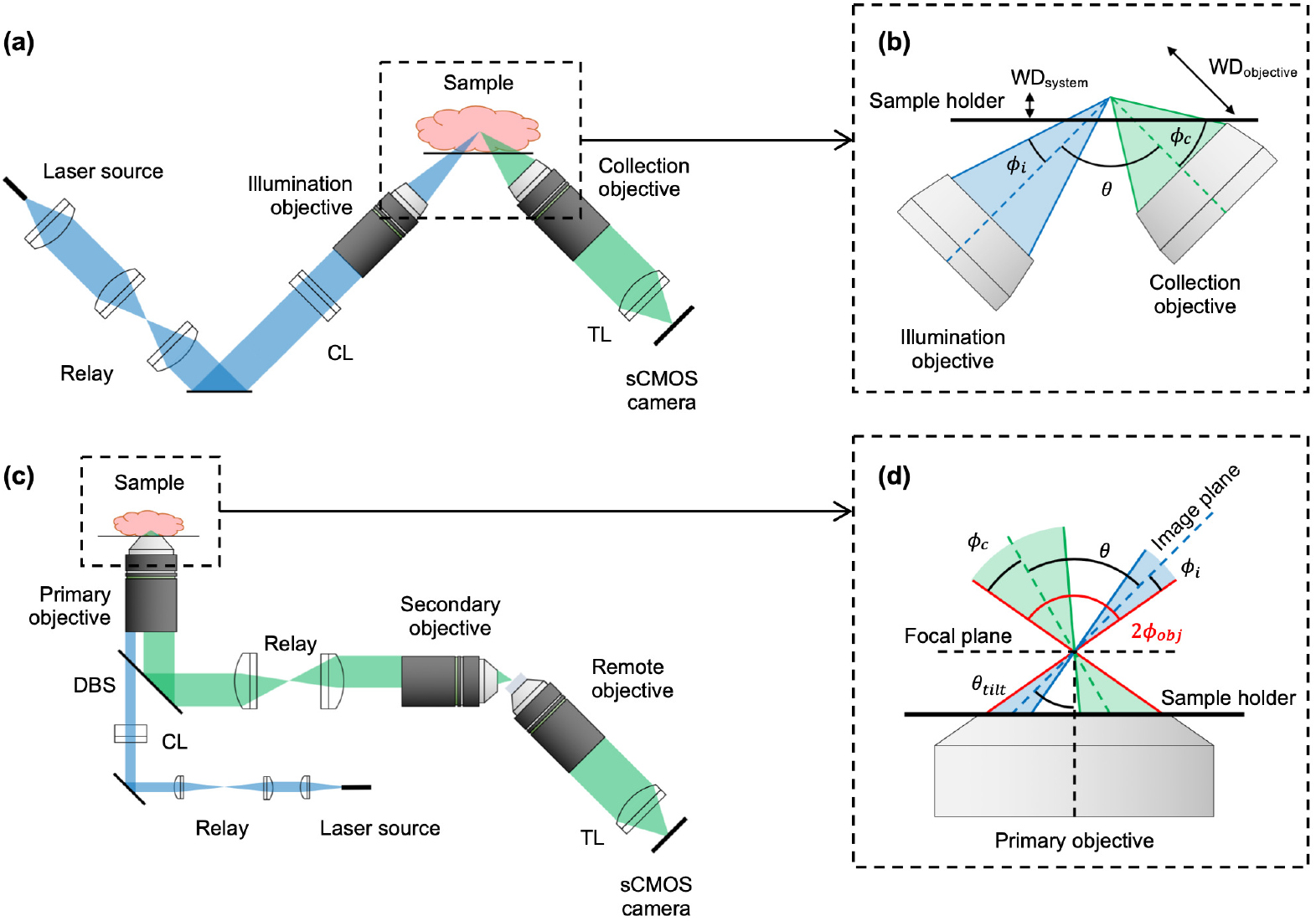
Overview of single- and dual-objective light-sheet microscope (LSM) architectures. (a) Optical schematic of a conventional orthogonal dual-objective (ODO) open-top light-sheet (OTLS) system, showing the illumination (blue) and collection (green) light paths. (b) Inset of illumination and collection objectives of an ODO system, showing that the system’s effective working distance (WD_system_) is much less than the collection objective’s working distance (WD_objective_). (c) Optical schematic of a non-orthogonal single-objective (NOSO) OTLS system, showing the illumination (blue) and collection (green) light paths. (d) Inset of the shared primary objective of a NOSO system, showing the angled light sheet and collection path. CL: cylindrical lens, TL: tube lens, DBS: dichroic beam splitter.

Single-objective light-sheet microscopes have recently been developed [15–21], which use a portion of an objective’s NA for illumination and a (typically larger) portion of the NA for collection (Fig. 1c). The use of a single objective allows the full working distance of the objective to be made accessible for imaging the sample. In addition, orienting the objective in the normal direction with respect to a horizontal sample holder leverages the ideal aberration-correction properties that have been meticulously engineered into high-quality microscope objectives when imaging through flat interfaces, thus relaxing sample index-matching requirements. Furthermore, since the conjugate planes of the illumination and collection beams are the same, scanning the beams (e.g. with a mirror) in tandem, while maintaining their alignment within the sample, can be simplified.

A significant challenge in implementing a single-objective light-sheet design is that when both light paths share the same objective, the illumination light sheet cannot be oriented orthogonally to the objective (Fig. 1d). This makes it challenging to efficiently capture an in-focus image of the fluorescence generated by the light sheet on a flat detector array (camera chip). An innovation that has made single-objective LSM possible is creating a remote focus with a secondary objective such that a tilted third objective can be used to image the remote light sheet onto a sCMOS detector array in the ideal orthogonal direction [15–21]. The details of this approach are described in Appendix A.

In light of the numerous LSM architectures and variations that have been developed in recent years, there is a natural desire amongst optical engineers to analyze their performance tradeoffs. While some analysis has been performed for a narrow subset of LSM configurations, such as examining different illumination profiles for dual-objective systems [22] and simulating various single-objective designs [23], a systematic comparison of diverse dual-objective and single-objective LSM designs has not been performed. Here we seek to fill this gap by presenting a quantitative analysis of dual- and single-objective LSM performance in response to several key design parameters. By quantifying design tradeoffs between different configurations, there is potential to identify “unreached” design spaces, which in turn will motivate future areas of innovation. For example, we describe a new non-orthogonal dual-objective (NODO) configuration that provides some advantages over orthogonal dual-objective (ODO) and non-orthogonal single-objective (NOSO) configurations for moderate-resolution applications.

It is impossible to provide a comprehensive guide for LSM design given the vast range of factors to consider such as sample characteristics (e.g. size, quantity, degree of optical clarity) and biological considerations (e.g. dynamic vs. static imaging, light dose), among others. Here we have focused our analysis on OTLS microscopy configurations for imaging fixed tissues that are optically cleared, in which optical scattering and refractive aberrations are assumed to be negligible. We also focus our analysis to assume conventional Gaussian illumination, which is the most-popular illumination method for LSM (note that a recent analysis of various LSM illumination schemes was performed by Remacha et al. [22]). With this context in mind, we examine a set of major performance metrics - primarily resolution and contrast - as functions of a specific set of design parameters: illumination and collection beam NAs and crossing angle, as constrained by currently available objectives. Certain imaging applications could warrant consideration of alternative parameters. Optical scattering, for example, is a key consideration for imaging live, uncleared specimens, and has been examined with Monte-Carlo simulations in the past for LSM [24] and dual-axis confocal microscopy [25,26]. Nonetheless, we believe that our work represents a valuable first step towards guiding the design of OTLS systems.

## 2. Analysis approach

The overall analysis approach we followed is shown in Fig. 2. We identified three key design parameters that, in the context of cleared-tissue imaging, are primary drivers of a system’s optical performance regardless of the specific architecture (single- or dual-objective) used: the NA of the illumination beam, the NA of the collection beam, and the angle between the two beams. Note that we make several assumptions for the sake of simplifying our analysis. We assume single-photon (linear) Gaussian illumination, which is used in the majority of LSM applications. In particular, pure Gaussian illumination is assumed, which is a fair approximation for real Gaussian beams that are truncated (apodized) beyond the 1/*e*^2^ intensity points. Further, we assume that the collection NA is larger than the illumination NA, which is true in nearly all LSM systems. This allows for straightforward mathematical descriptions of geometric constraints, which in turn limit the numerical range of the design parameters that are analyzed (see next section). We assess optical performance in terms of spatial resolution and contrast (as determined by a simulated fluorescent bead phantom).

**Fig. 2.**
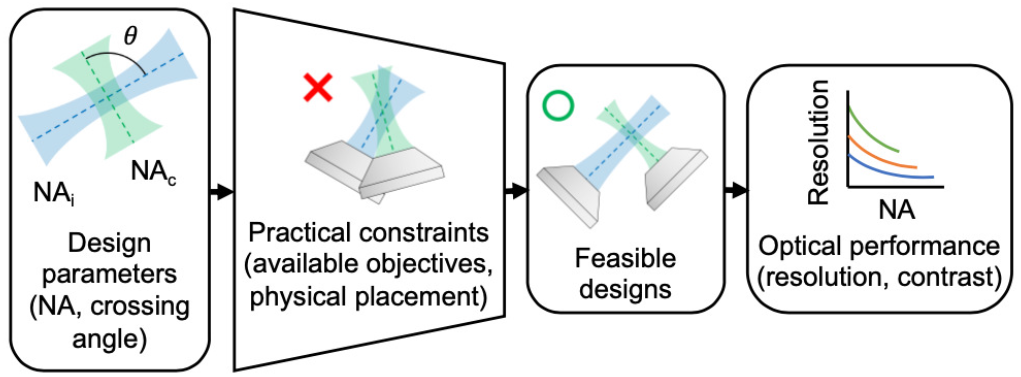
Outline of the analysis approach followed. Three key design parameters were identified that, in the context of cleared tissue imaging, primarily determine the optical performance of an LSM system (regardless of specific architecture): the numerical aperture (NA) of the illumination beam, the NA of the collection beam, and the crossing angle between them. We then introduced limits on these parameters based on practical constraints including available objectives and the physical placement of those objectives. These limits give rise to a range of feasible designs. Finally, we assess the optical performance of these designs in terms of spatial resolution and contrast.

### 2.1. Geometric constraints and simplifying assumptions for dual-objective systems

The optical path for an ODO system is shown in Fig. 1a. Note that these systems generally include scanning mirrors to translate the light sheet and/or stages to scan the specimen, but these have been omitted for simplicity in both Fig. 1a and Fig. 1c. For optically clear tissue specimens, unrestricted lateral sample extent is generally preferred. As such, we assume an OTLS architecture for both single- and dual-objective systems, which allows for a more-consistent comparison between various LSM architectures.

While the two objectives in a dual-objective system are typically placed orthogonally to one another, in principle their crossing angle is not restricted to90°. We assume that the objective housing does not occupy additional space beyond the optical cone (NA) of the objective. As such, the placement of the objectives is restricted by the crossing angle and the cone angles of the individual beams, as shown in Fig. 1b. The half cone angle for each objective is determined by:

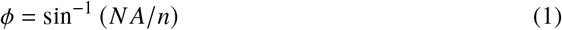

Here, *ϕ* is the half cone angle, *NA* is the numerical aperture of the objective, and *n* is the refractive index of the immersion medium (assumed to be water, n = 1.33, for this analysis). The objectives cannot physically collide, so the minimum crossing angle permitted in this system is the angle at which the illumination and collection cones begin to intersect:

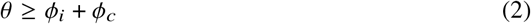

Here, *θ* is the crossing angle between the optical axes of the two objectives, while *ϕ_i_* and *ϕ_c_* are the half cone angles of the illumination and collection objectives, respectively. In order to maintain an OTLS geometry, the objectives also cannot cross the horizontal sample holder as they would collide with the sample. The maximum crossing angle physically allowed is then constrained by:

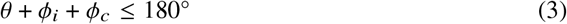

This relationship assumes that the thickness of the sample holder and sample itself are negligible. Note that a crossing angle larger than 90° would never be used in practice. Equations (2) and (3) provide physical limits on how a dual objective system can be constructed, which constrain the numerical range of our simulations. Importantly, the observation that the crossing angle *θ* does not need to be set to 90° (as is the case in conventional orthogonal configurations) gives rise to a novel NODO design that we will examine closely later.

### 2.2. sGeometric constraints and assumptions for single-objective systems

Figure 1c shows the light path of a generic NOSO LSM system (with the objective arranged below the sample as previously described). As shown in Fig. 1d, this configuration results in a light sheet that is tilted at some angle *θ_tilt_* relative to the optical axis of the objective. The crossing angle of the illumination and collection cones is constrained by the physical NA of the shared primary objective. Assuming that the crossing angle is maximized for a given illumination and collection NA (which offers the best performance, as shown later), the crossing angle θ is given by:

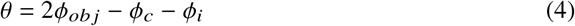

Here, *ϕ_obj_*, *ϕ_c_*, and *ϕ_i_* are the geometric half cone angles of the shared primary objective, the collection beam, and the illumination beam, respectively. Note that as in the dual-objective analysis, these geometric angles are assumed to be within a high-index medium (water, *n* = 1.33, in this analysis).

Additional geometric analyses of the NOSO architecture (used to limit our simulations), and of light behavior at the remote focus, are provided in Appendix A. In particular, we assume that techniques are employed (as described by others [19–21]) to ensure that the collection beam can be imaged by a high-NA remote objective (and detected by the camera) with 100% throughput and negligible aberration.

### 2.3. Non-orthogonal dual-objective design

As mentioned previously, our analyses will include a novel NODO architecture, which combines attractive elements of both the ODO and NOSO configurations. In a NODO configuration, the collection objective is kept perpendicular to the sample holder and a second low-NA objective is used for non-orthogonal illumination (Fig. 3). Similar to NOSO systems, this offers the ability to access the full working distance of the collection objective for sample imaging, and significantly reduces sensitivity to index mismatch. Note that the latter issue (index mismatch) can be a major source of aberrations in ODO systems, where high-NA beams must transition between different media at highly oblique angles.

**Fig. 3.**
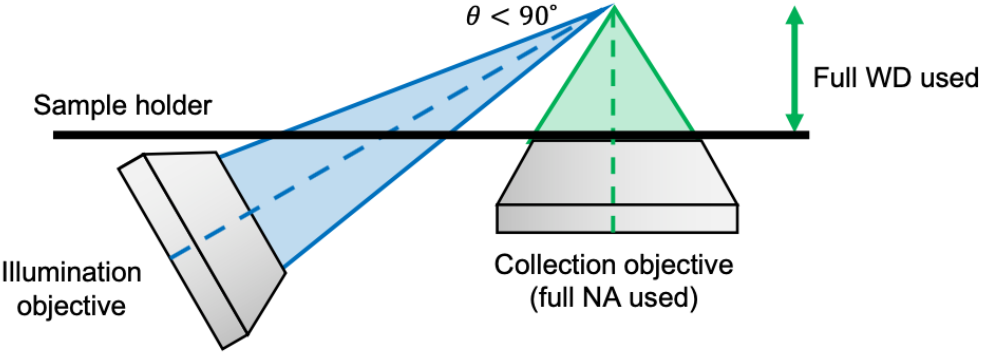
Geometry of a proposed non-orthogonal, dual-objective (NODO) design. The crossing angle θ is less than 90° so that the collection objective can be oriented in the normal direction with respect to the horizontal sample holder, as in a NOSO system. This allows the full working distance of the collection objective to be used and minimizes sensitivity to refractive-index mismatch. Unlike the NOSO architecture, NODO also allows the full NA of the collection objective to be used and allows for a higher crossing angle between the illumination and collection beams (with implications for axial resolution and contrast).

Compared with NOSO designs, a major advantage of the NODO configuration is that since the illumination and collection beams do not need to share a single objective, there is more design flexibility in terms of beam NAs and crossing angle. One downside to a NODO system is that the illumination and collection paths no longer share a conjugate plane as in a NOSO system. This makes rapid scanning and descanning of these beams more challenging, such that the system may be less ideal for live-sample imaging. Additionally, since the collection objective is oriented normal to the sample holder, the angled illumination objective must have a relatively low NA and a long-enough working distance to be positioned adjacent to the collection objective. Nonetheless, the practical advantages of the NODO configuration for cleared-tissue imaging warrant further consideration and quantitative comparison with conventional ODO and NOSO systems.

### 2.4. Quantitative output metrics

In cleared-tissue imaging, image quality is the primary performance metric of interest. Volumetric imaging speed is of concern, but typically scales in a predictable way with spatial resolution if samples are sufficiently labeled and photon counts are not a limiting factor. Light dose (i.e. photobleaching and photodamage) is not as critical compared with live-tissue imaging applications. We thus quantified the performance of different configurations in terms of the spatial resolution and contrast that is theoretically possible for a given system. We computed axial and lateral resolution values based on the simulated point spread function (PSF) of each system. We analyzed the contrast of each system by simulating the system’s response to a virtual fluorescent bead phantom. Additional details regarding these resolution and contrast computations are provided in Appendix B.

## 3. Analysis results

In this study, we used the geometric constraints identified above to identify key input parameters for each microscope configuration. For dual-objective systems, the input parameters are: illumination NA, collection NA, and crossing angle (where ODO systems have a crossing angle of 90° and NODO systems have a crossing angle less than 90°). In single-objective systems, the input parameters are: primary objective NA, effective illumination NA, and effective collection NA. Note that these three parameters uniquely determine the crossing angle of the beams, as expressed in Eq. (4). We limited our simulations to a range of primary objective NAs (i.e. collection-objective NA for dual-objective systems) ranging from 0.4 to 1.0 based on currently available commercial objectives that are suitable for imaging cleared tissues (Appendix C). Axial resolution, lateral resolution, and contrast were computed as functions of the listed design parameters. Rather than presenting every possible combination of input parameters and output metrics, we focus our presentation on a number of observations that are deemed to be most critical for designers.

### 3.1. Resolution

#### 3.1.1. For dual-objective systems, larger crossing angles improve axial resolution until ~60°

We first consider how the axial resolution of a conventional ODO system compares to a NODO system by examining how it varies as a function of crossing angle. Figure 4a shows this relationship for two pairs of illumination and collection NAs (*NA_i_* = 0.1, *NA_c_* = 0.4 and *NA_i_* = 0.2, *NA_c_* = 0.7). As expected, axial resolution improves with crossing angle, with the ODO configuration thus providing the best axial resolution. However, the gains in axial resolution are minimal beyond angles of ~60°, a finding that is consistent across NA pairs beyond those shown. Conveniently, this angle is readily achievable (in terms of physical placement of objectives) for most choices of illumination and collection NA, including for the NODO case in which the collection objective is oriented normal to the horizontal sample holder. Therefore, we use 60° as the crossing angle for all further analyses of NODO systems.

**Fig. 4.**
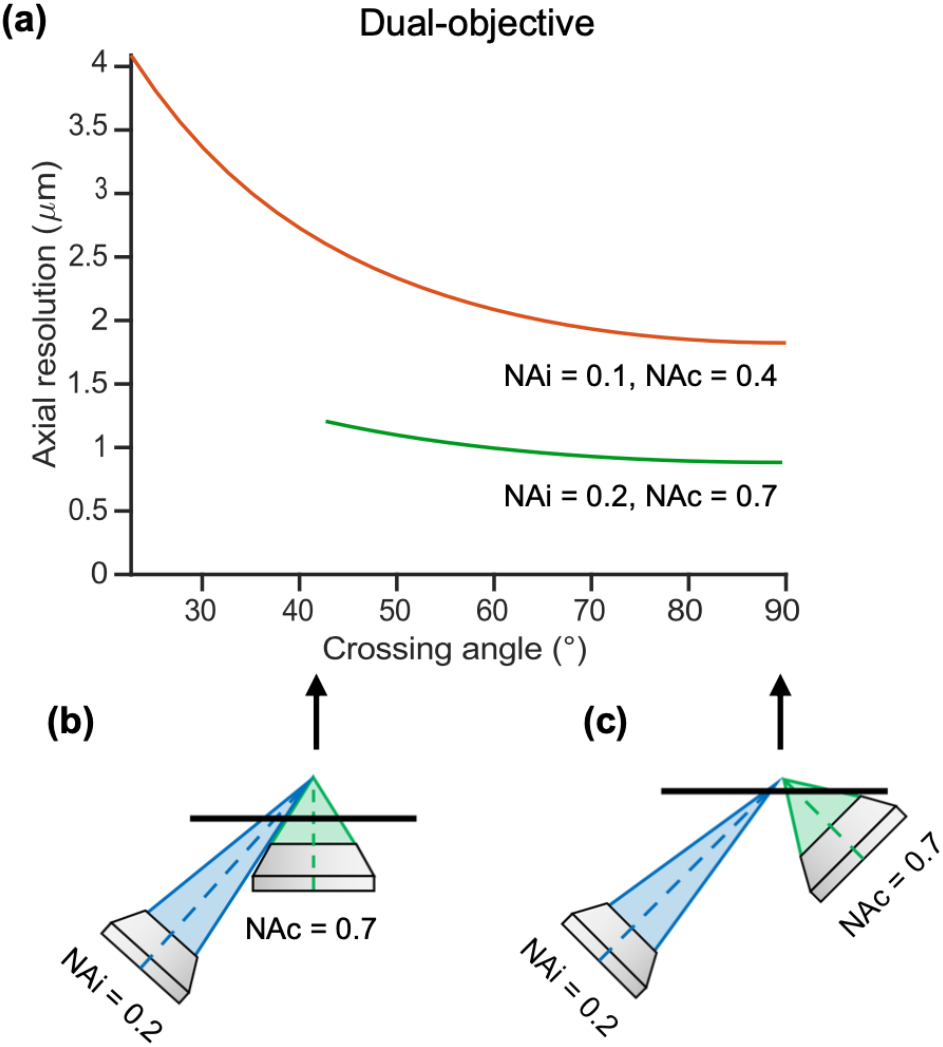
Axial resolution of a dual-objective system. (a) Axial resolution depends strongly on crossing angle at small angles but does not improve much beyond angles of 60°. Curves for two example illumination and collection NA pairs are shown (*NA_i_* = 0.1, *NA_c_* = 0.4; *NA_i_* = 0.2, *NA_c_* = 0.7), but these trends are consistent across different choices of illumination and collection NA. (b) The minimum angle plotted for each system is the minimum crossing angle before the objective cones overlap. (c) The maximum crossing angle is 90° (the conventional ODO case), which yields the maximum possible axial resolution.

#### 3.1.2. For all systems, axial resolution is highly dependent upon illumination NA

Besides crossing angle, a primary driver of axial resolution is the light-sheet thickness as determined by the illumination NA. Figure 5 shows the impact of illumination NA on axial resolution for each system architecture. There are diminishing returns as the illumination NA is increased (beyond ~0.3). Two collection NAs are shown in each plot (0.4 and 0.7). This trend is similar for other collection NA choices with a slightly diminishing impact of illumination NA on axial resolution as collection NA is increased. This is because as collection NA increases, the limited depth of focus of the collection objective increasingly provides optical sectioning, thereby reducing the relative contribution of light-sheet thickness to axial resolution. An implication of these observations is that while isotropic resolution at high NAs may be desirable in theory [5, 27–30], the simplicity and geometric advantages of using a relatively low illumination NA (~0.3) may offer an ideal compromise for many practical applications of LSM.

**Fig. 5.**
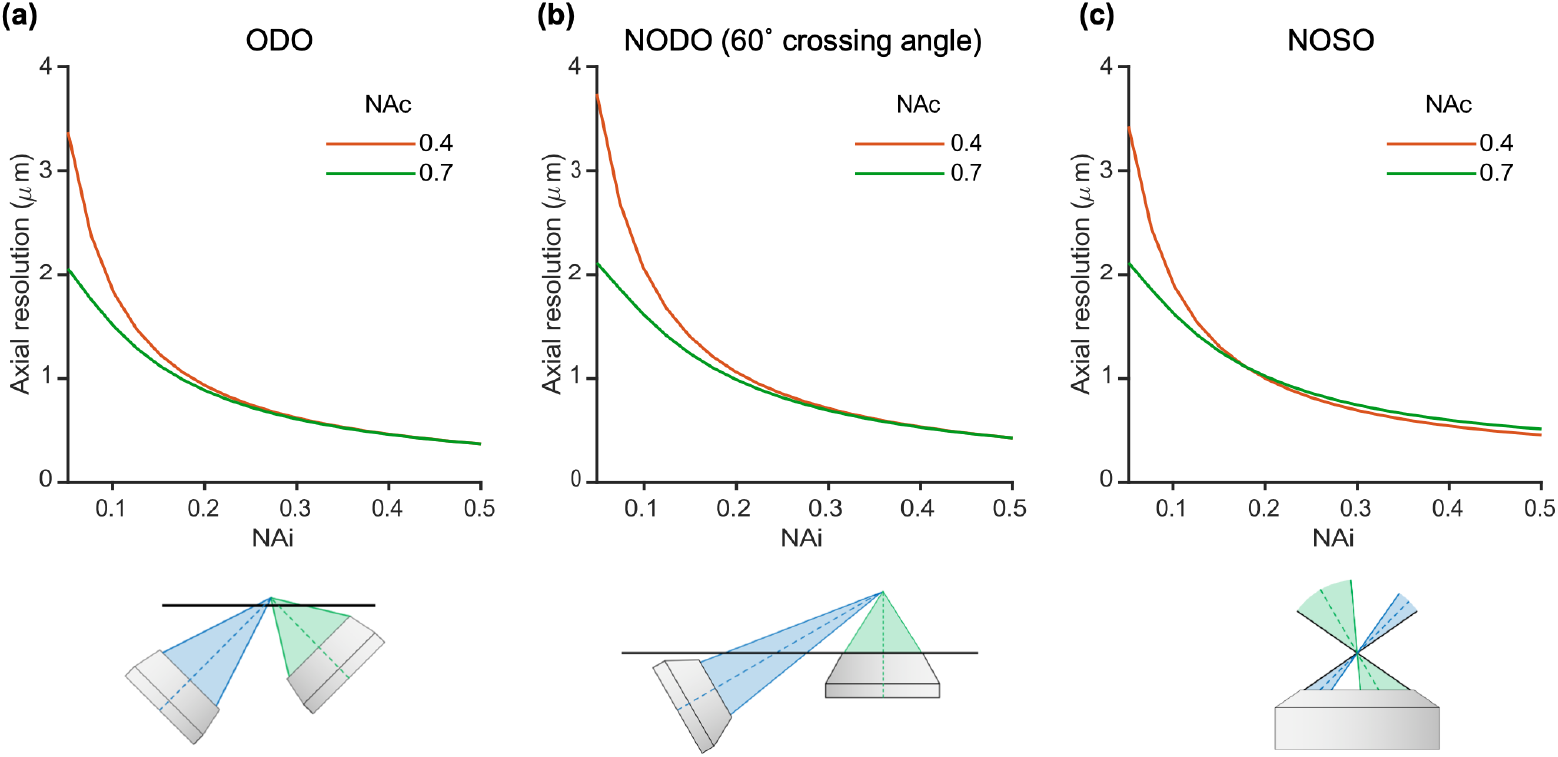
Impact of illumination NA on axial resolution for (a) an ODO system, (b) a NODO system at a crossing angle of 60°, and (c) a NOSO system using a 1.0 NA primary objective. In all cases, illumination NA is a primary driver of axial resolution. Collection NAs of 0.4 and 0.7 are shown, but trends are similar for other collection NAs. A diagram of each configuration is shown below each corresponding plot.

#### 3.1.3. For dual-objective systems, lateral resolution is entirely determined by collection NA

The previous plots showed that in a dual-objective system, the axial resolution is primarily determined by illumination NA and crossing angle. We will now consider which factors are most impactful to lateral resolution in a dual-objective system. Figure 6 shows how lateral resolution varies as a function of collection NA for an ODO system (a) and a NODO system with a 60° crossing angle (b). In these plots, the different colors correspond to different choices of illumination NA. In all cases, increasing the collection NA improves the lateral resolution. We also see that the different curves in each plot almost completely overlap, indicating that lateral resolution is not influenced greatly by illumination NA. Further, the curves are nearly identical between the ODO and NODO plots, indicating that lateral resolution is also independent of crossing angle. In summary, lateral resolution in a dual-objective system is almost entirely determined by collection NA. Combined with the previous finding that axial resolution in a dual-objective system is primarily determined by illumination NA and crossing angle, we observe that using two objectives, which allows axial and lateral resolution to be decoupled, provides a designer with significant flexibility for optimizing system performance.

**Fig. 6.**
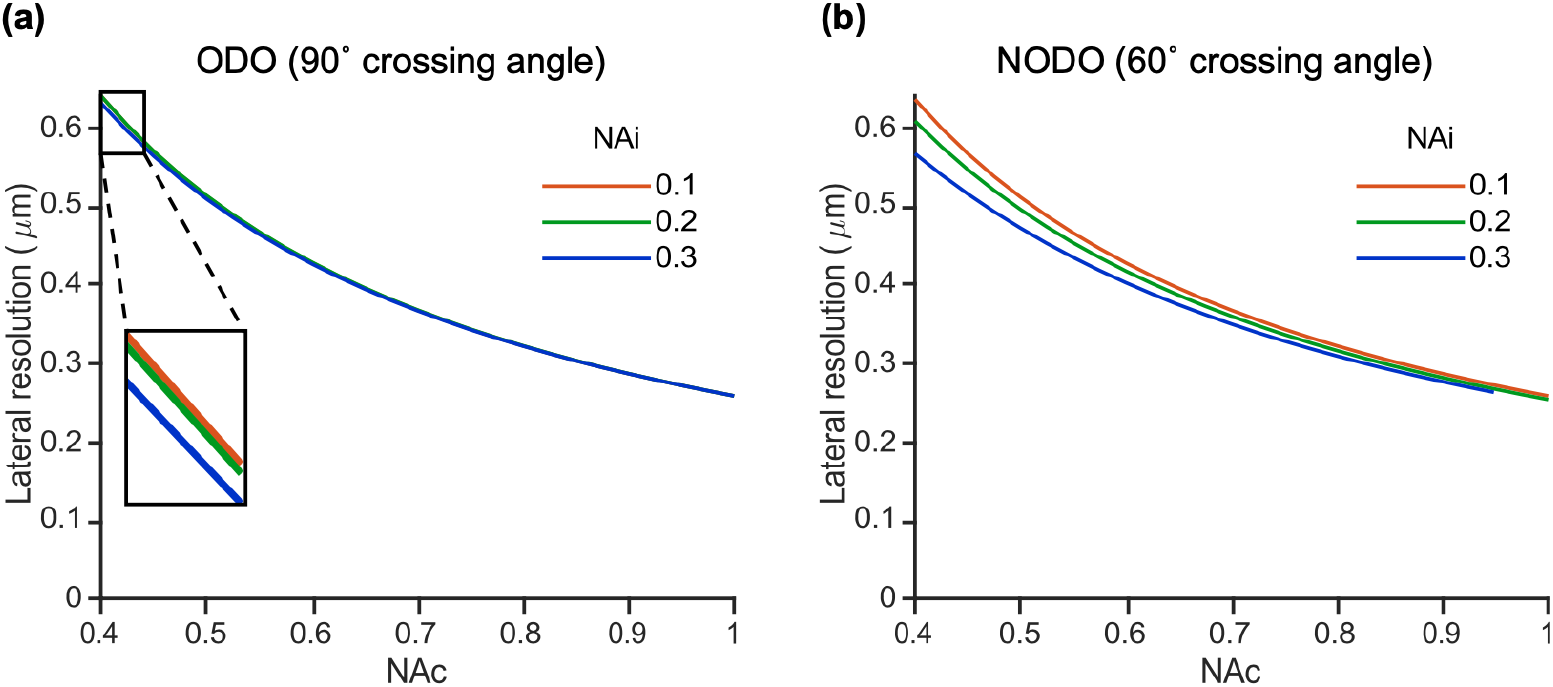
Lateral resolution as a function of collection NA for (a) an ODO system and (b) a NODO system with a 60° crossing angle. Different curves on each plot indicate different choices of illumination NA. Curves are largely unchanged both within each plot and between plots, indicating that lateral resolution in a dual-objective system is primarily determined by collection NA and is independent of illumination NA and crossing angle.

#### 3.1.4. For single-objective systems with moderate NA, axial and lateral resolution must trade off

In contrast to the decoupling of axial and lateral resolution in a dual-objective system, axial and lateral resolution in a single-objective system are intrinsically linked because the two light paths share a single objective. Figure 7 shows two example NOSO systems: one using a moderate-NA primary objective (0.75, a) and one using a high-NA primary objective (1.0, b). In each plot, the primary objective NA is held constant, showing the different performance combinations a designer could achieve with a given optical element. For simplicity, the illumination NA is also held constant (selected as 0.2 for these examples) as it is generally restricted to a low value in a NA-constrained NOSO system. The plots show how axial and lateral resolution vary as functions of collection NA. Note that varying the collection NA implicitly changes the effective crossing angle of the illumination and collection light paths since they share one objective (as seen in Eq. (4)).

**Fig. 7.**
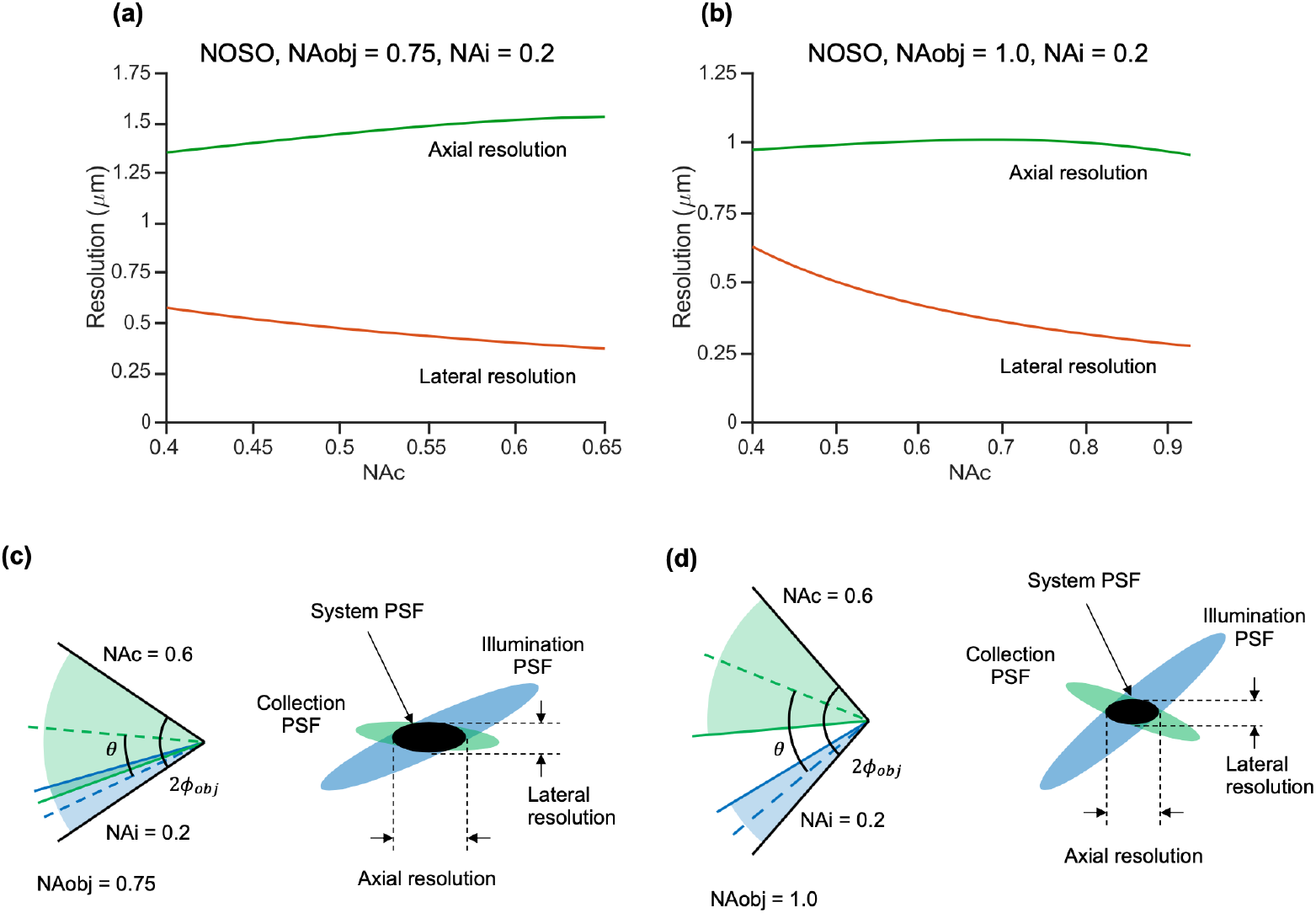
Lateral and axial resolution as a function of collection NA for a NOSO system in the case of (a) a moderate-NA primary objective (0.75 NA) and (b) a high-NA primary objective (1.0 NA). In each plot, the primary objective NA is held constant to reveal the different performance combinations a designer could achieve with a given optical element. For simplicity, the illumination NA is also held constant (0.2). Note that changing the collection NA implicitly changes the effective crossing angle in our NOSO simulations. (c) Cone angles and PSF schematic for a moderate-NA shared primary objective. At moderate objective NAs, the crossing angle θ is constrained to relatively small values. Any further reduction in crossing angle, resulting from an increase in collection NA, leads to a degradation in axial resolution. Thus, there is a tradeoff between axial and lateral resolution. (d) Cone angles and PSF schematic for a high-NA shared primary objective. The crossing angle θ is relatively large in all cases, such that axial resolution is less sensitive to changes in collection NA. For ease of comparison, the same collection NA (0.6) is shown in both (c) and (d).

We first consider the moderate-NA case (0.75 NA shared primary objective). Unsurprisingly, higher collection NAs result in superior lateral resolution. Interestingly, the increase in collection NA causes a degradation in axial resolution because it results in a smaller crossing angle between the illumination and collection light paths (Fig. 7c), which lengthens the axial dimension of the system PSF. This finding highlights the implicit tradeoff between axial and lateral resolution in a single objective system.

At higher NAs (1.0 NA shared primary objective), the crossing angle remains relatively high regardless of effective collection NA. Therefore, axial resolution is not strongly dependent upon collection NA. In other words, for a high-NA NOSO system, axial and lateral resolution do not trade off to the extent that they must for a NOSO system with a lower primary objective NA. However, the beams must still share an objective, restricting the overall maximum collection NA and crossing angle that can be achieved in comparison to a dual-objective system.

### 3.2. Contrast

#### 3.2.1. For dual-objective systems, larger crossing angles improve contrast until ~60°

In addition to the insights into resolution that have been discussed so far, we also identified quantitative trends regarding the contrast of single- and dual-objective systems. First, we consider the contrast of a dual-objective system as a function of crossing angle. Figure 8a shows how contrast (analyzed with a simulated fluorescent bead phantom as described in Appendix B) of a dual-objective system varies as a function of crossing angle. Different curves represent different illumination NAs. A collection NA of 0.7 is used in this example, though the trends are similar for other collection NAs (with slight improvements in contrast at higher collection NAs). For example, plots for collection NAs of 0.4 and 1.0 are provided in Appendix D. Like in Fig. 4, the minimum and maximum crossing-angle cases are depicted in Fig. 8b and Fig. 8c, respectively. Contrast improves at higher crossing angles as the collection and illumination light paths are more spatially separated. As with axial resolution of a dual-objective system, this dependence is strongest at small angles, and contrast improves little beyond 60°. This reinforces the observation that a NODO system with a ~60° crossing angle is ideal, with optical performance close to that of an ODO system.

**Fig. 8.**
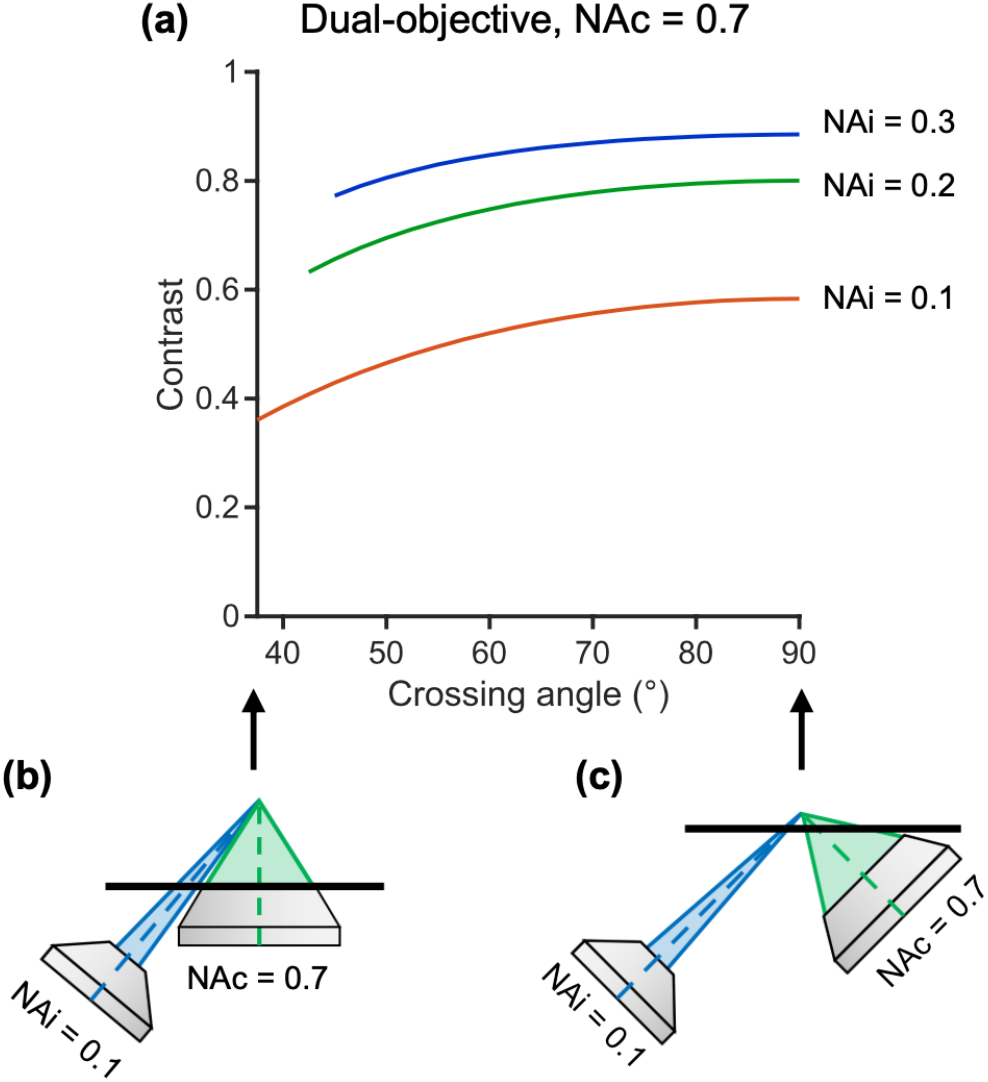
Contrast of a dual-objective system. (a) Impact of crossing angle on contrast is plotted for three illumination NAs (different curves). A collection NA of 0.7 is used in this example, though the trends shown hold for other collection NAs (with slight improvements in contrast overall at higher collection NAs). The minimum angle plotted for each system is the minimum crossing angle before the illumination and collection cones overlap. The maximum angle plotted is 90° (the conventional ODO case) which gives the maximum possible contrast. These cases are shown for a 0.1 illumination NA and 0.7 collection NA system in (b) and (c), respectively. Contrast improves with higher crossing angles, but gains begin to saturate around 60°.

#### 3.2.2. Contrast improves with higher illumination NA, and dual-objective systems exhibit improved contrast over single-objective systems at moderate NAs

Figure 9 compares the contrast of an ODO, NODO (60° crossing angle), and NOSO system as a function of illumination NA. In order to provide a fair comparison, we consider a moderate-NA case (0.7, a) and high-NA case (1.0, b), keeping the collection/primary objective NA constant for each case. In other words, the plot shows the different configurations that could be constructed with the same optical element (a 0.7 or 1.0 NA objective). For simplicity, the effective collection NA is held constant (0.5 and 0.7 in a and b, respectively) for each NOSO curve in Fig. 9. Different choices of collection NA change the upper limit on illumination NAs (as the beams must share one objective) but otherwise result in similar trends. For all systems, contrast improves with illumination NA but offers diminishing returns at higher illumination NAs (similar to axial resolution, Fig. 5). In addition, the ODO and NODO systems exhibit superior contrast to comparable NOSO systems at moderate NA. The main reason for this is that crossing angle plays a significant role in determining contrast, and dual-objective systems can achieve higher crossing angles, particularly in the moderate-NA case. At high NA, this effect is less significant, and all three systems provide similar contrast.

**Fig. 9.**
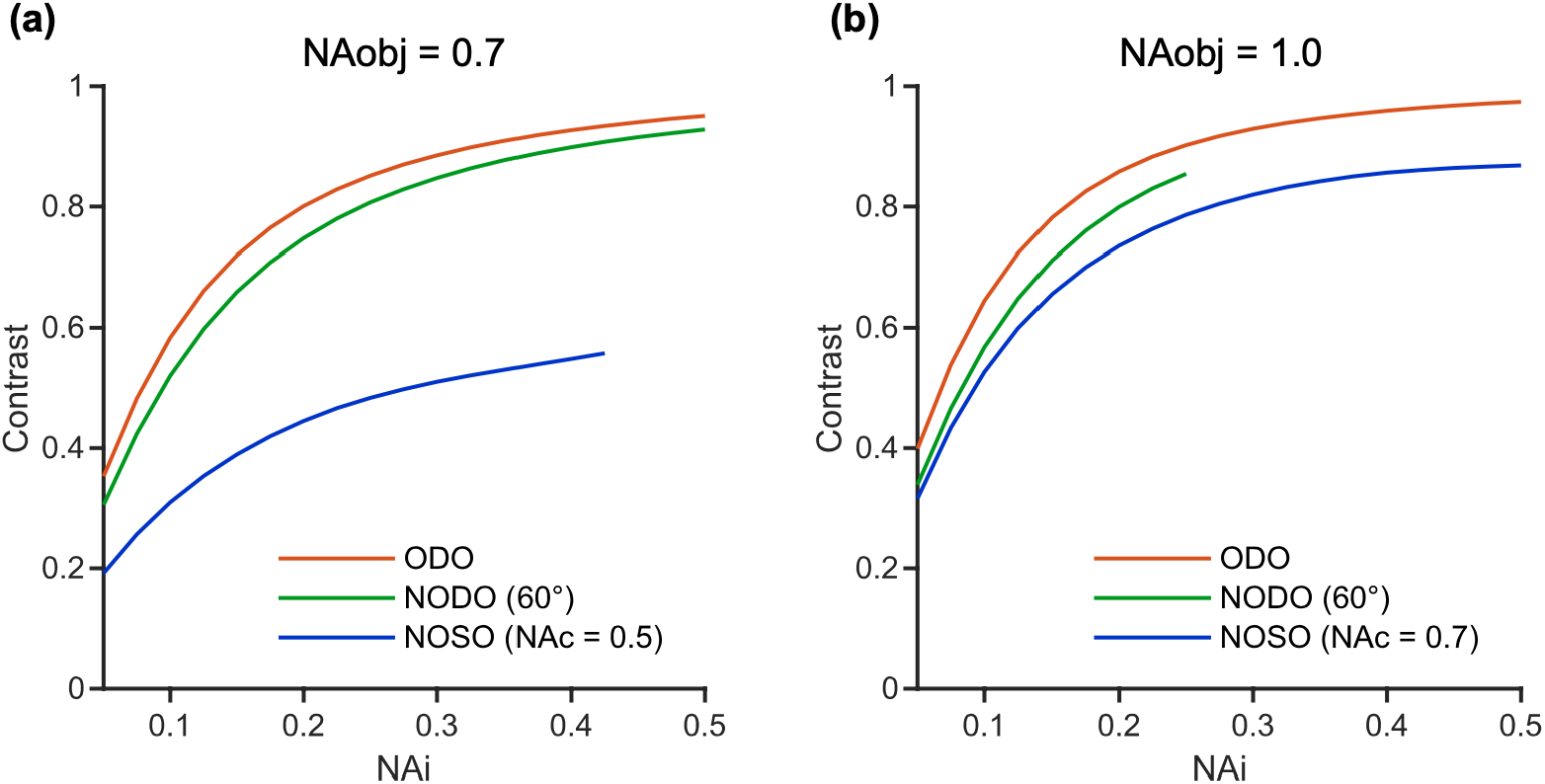
Comparison of contrast for ODO, NODO (60° crossing angle), and NOSO systems that can be built using (a) a moderate-NA (0.7) primary objective and (b) a high-NA (1.0) primary objective (i.e. the collection objective for dual-objective systems or the shared primary objective for single-objective systems). For simplicity, the effective collection NA is held constant for each NOSO curve (0.5 for the moderate-NA case and 0.7 for the high-NA case). For all systems, increasing the illumination NA improves contrast but offers diminishing returns at higher illumination NAs. The NOSO system provides poorer contrast than the ODO or NODO systems in the moderate-NA case, while all systems provide similar contrast in the high-NA case.

## 4. Discussion

We have identified several key trends in terms of the resolution and contrast of single- and dual-objective LSM systems for cleared-tissue imaging, which we believe will be valuable as a general guide for system designers. In particular, we evaluated three OTLS microscope architectures: a conventional ODO system, a conventional NOSO system, and a novel NODO system.

Regarding the resolution of these systems, there are four main observations.

1. For dual-objective (ODO and NODO) systems, larger crossing angles improve axial resolution until around 60° (Fig. 4).
2. For all systems, axial resolution is significantly affected by illumination NA, with diminishing returns at higher illumination NAs (Fig. 5).
3. For dual-objective systems, lateral resolution is entirely determined by collection NA, meaning that axial and lateral resolution are decoupled from one another (Fig. 6).
4. For single-objective (NOSO) systems at moderate NA, axial and lateral resolution must trade off (Fig. 7).

We also identified two trends regarding system contrast.

1. For dual-objective systems, larger crossing angles improve contrast until around 60° (Fig. 8).
2. Contrast for all systems improves with illumination NA, with diminishing returns at the highest illumination NAs, and dual-objective systems exhibit better contrast than single-objective systems when the same moderate-NA objective is used (Fig. 9).

Collectively, these observations indicate that rather than one design being universally superior to another, there are specific benefits and drawbacks to each configuration depending upon the application.

There are a number of pros and cons for a NODO system. From a practical standpoint, a NODO system allows the full working distance and full NA of a collection objective to be utilized. The collection objective should ideally be oriented normal to a horizontal sample holder/interface, where refractive aberrations and index-matching requirements are minimized with modern well-corrected objectives. However, similar to a NOSO system, NODO requires a strategy to image a highly oblique non-orthogonal light sheet onto a camera chip (typically flat), such as by re-imaging a tilted remote focus. This adds a degree of optical complexity absent from an ODO system (though the remote focus does not limit performance to the extent it does in a NOSO system). Rapid time-lapse imaging of a localized 3D volume is also more challenging in a NODO system than in a NOSO system, making it less ideal for dynamic live-tissue imaging applications. Additionally, the use of multiple objectives means that, unlike a NOSO system, a NODO system cannot be easily integrated into a commercial widefield or confocal microscope.

Our quantitative analysis reveals several important aspects of NODO optical performance. For example, the NODO system (with ~60° crossing angle) provides axial resolution comparable to an ODO system. Like an ODO system, the NODO system would require matched illumination and collection objectives to achieve isotropic resolution. However, using a relatively low illumination NA (~0.3) offers similar resolution and contrast performance (Fig. 5 and Fig. 9) but with the simplicity and geometric flexibility that a lower-NA illumination arm can offer, which may be most ideal for many applications. Image contrast for a NODO system is also comparable to an ODO system and better than a NOSO system. In addition, the NODO architecture decouples axial and lateral resolution, giving the designer more flexibility than a NOSO system.

It is important to note that the NODO design described here has some mechanical similarities to light sheet theta microscopy (LSTM), with both designs featuring a perpendicular collection objective and angled, non-orthogonal light sheet(s) [8]. Instead of using a remote focus to bring an entire 2D image into focus at the camera, LSTM only captures fluorescence from the central row of pixels (at the beam waist of the light sheet) in a manner similar to line-scanned dual-axis confocal (LS-DAC) microscopy [24, 31, 32]. This improves axial resolution slightly, as only the absolute thinnest part of the light sheet is used for imaging, but requires complicated mechanical scanning in order to align and synchronize the focus of the light sheet with the focal plane of the detection objective and the rolling shutter of the camera. The LSTM design is also not as optically efficient compared to most LSM systems since most of the excited fluorescence is rejected by the sCMOS rolling shutter.

The advantages of the NODO architecture are most impactful for moderate-NA imaging systems. For low-NA systems, an ODO system is generally sufficient: long working-distance objectives and minimal off-axis aberrations mean that the complexities of a NODO architecture (e.g. having a tilted remote focus) are not warranted. At high NAs, a NOSO system allows the illumination and collection beams to have relatively large cone angles and crossing angles, which results in acceptable levels of resolution and contrast. However, as with any high-resolution (high-NA) system, the working distance and field of view of such a system will be limited, and imaging speeds will be slow. Finally, for moderate-NA systems (i.e. 0.5 < NA < 0.8), the NODO architecture offers a combination of imaging performance (resolution and contrast), working distance, and field of view that is challenging to achieve with other architectures.

In summary, we have performed a quantitative analysis of how several key design parameters affect the performance of various OTLS microscope architectures. Our analysis reveals regimes in which various architectures are most ideal, and also shows that a NODO configuration may offer benefits for addressing cleared-tissue imaging applications where moderate spatial resolution is desired.

### Appendix A: Remote focus geometric analysis and NA limits for single-objective systems

When a sample is illuminated with a non-orthogonal light sheet in a single-objective light-sheet microscopy (LSM) system, the resultant image is no longer perpendicular to the optical axis of the system. In order to form an in-focus image at the camera, a remote focus can be used to optically rectify the image at the camera plane (Fig. 10a) [33]. Dunsby employed this method to develop the first single-objective LSM system by combining several precursor designs [15, 34, 35].

From Fig. 10b, we see that the angle at which the light sheet is tilted relative to the optical axis in a single-objective system, *θ_tilt_*, can be expressed as:

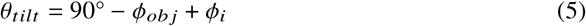

**Fig. 10.**
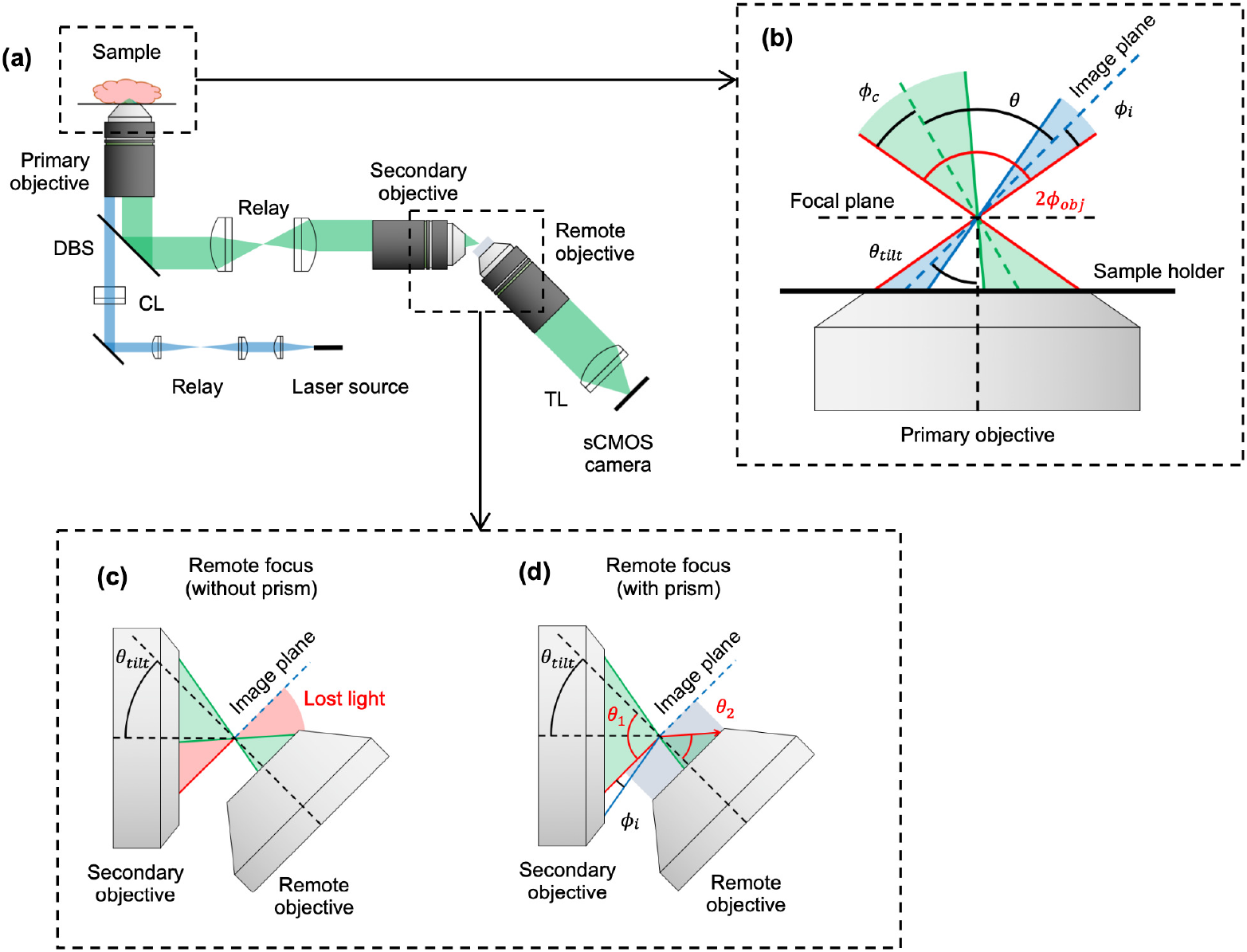
(a) Optical schematic of a non-orthogonal single-objective (NOSO) LSM system, showing the illumination (blue) and collection (green) light paths. (b) Inset of the shared primary objective of a NOSO system showing the angled light sheet and collection path. (c,d) Inset of the remote focus of a NOSO system (c) without a refractive prism, showing some light is lost at the remote focus and (d) with a refractive prism, showing collection of otherwise lost light. CL: cylindrical lens, TL: tube lens, DBS: dichroic beam splitter.

Here, *ϕ_obj_* and *ϕ_i_* are the geometric half cone angles of the shared primary objective and the illumination cone, respectively.

Consequently, the image formed by the light-sheet-generated fluorescence is also tilted, which prevents an in-focus image from being formed on an untilted camera. An intermediate image plane is created along the collection light path using a secondary objective, essentially creating an exact 3D replica of the angled fluorescence sheet. The virtual light sheet is then imaged onto the camera by a final “remote” objective, which is physically tilted by *θ_tilt_* such that it is normal to the light sheet, thus bringing the light sheet into focus at the camera chip (Fig. 10c). Bouchard et al. subsequently presented another single-objective LSM system, and was among several groups to present variations on Dunsby’s work that laterally (rather than axially) scanned the light sheet [16–18].

While the systems described above allow an in-focus image to be generated from a single-objective, the tilt of the remote objective often causes a portion of the light (related to the degree of tilt) to fall outside the acceptance cone of the remote objective, thus reducing the effective collection numerical aperture (NA) of the system (Fig. 10c). To mitigate this NA loss, a refractive element can be placed at the image plane to refract light into the remote objective Fig. 10d. The element is arranged such that the intermediate image is formed on the surface of the element and such that this image is in the focal plane of the remote objective. This principle is described with a prism below (as presented by [19, 21]), but other methods such as a diffraction grating can also be used [20]. This development means that using a remote focus to correct for a tilted image plane does not by itself necessitate a reduction in collection NA (resolution and optical throughout).

To understand the extent to which a prism can mitigate NA loss, it is useful to consider the “limiting” incident angle in terms of rays that could theoretically be collected. If a prism of refractive index *n*_2_ is placed at the front of the remote objective, which is an immersion objective designed for index *n*_2_, then a theoretical ray incident at the limiting angle travels in the image plane (red arrow in Fig. 10c). This ray is incident on the prism at 90° relative to the surface normal. Applying Snell’s law to such a ray gives:

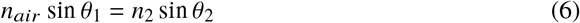

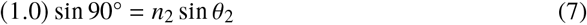

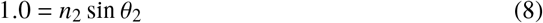

From this equation we can find the remote objective NA necessary to collect this ray:

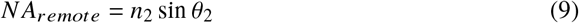

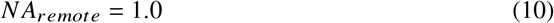

In theory, any remote objective with an NA of at least 1.0 can collect all rays incident at angles up to 90°, that is, all rays incident on the front surface of the prism. This is true regardless of the remote objective tilt. Any rays incident at angles larger than 90° hit the side of the prism rather than falling incident on the front surface of the prism and therefore can never be collected, regardless of the objective used. Because the ray at the limiting angle is also parallel to the central axis of the “virtual” illumination cone, this condition can be written mathematically as:

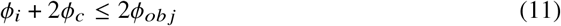

This gives a fundamental maximum on the collection NA that may be used for a given primary objective. Note that this limit arises from sharing a primary objective, not from the remote focus itself. Equation (11) together with Eq. (4) describe the physical limits on a single objective system that we use to limit our simulation conditions.

Practically speaking, light entering at such an extreme angle would suffer significant loss due to reflection (Fresnel) at the air/prism interface. The interface would also alter the polarization of the collected light, which may be undesirable. In practice, a designer would want to limit the collection light path to avoid such extreme angles. For simplicity, we will not consider these effects in our analysis, with the acknowledgment that this edge case represents a maximum limit on collection and illumination NA rather than a practical region of operation.

### Appendix B: Resolution and contrast computation

#### Spatial resolution computation

In order to quantify spatial resolution, we computed the intensity point spread function (PSF) of each microscope configuration in MATLAB (R2019b, MathWorks, Natick, MA, USA) using the adjoint method [24, 36–38]. For each simulation condition, 3D intensity PSFs were first computed for the illumination and collection beams. Gaussian illumination was assumed (at various NAs), along with uniform collection of angularly isotropic fluorescence emission (also at various NAs). The illumination PSF was rotated relative to the collection PSF at various crossing angles [16], and the overall system PSF (intensity) was calculated as the product of the two beams at each point in space (Fig. 11a). The system PSF describes the 3D spatial intensity distribution of signal that is detected from a point source of fluorescence. Note that due to the oblique crossing angle, in most cases the resultant system PSF is neither perfectly symmetric nor on-axis. The resolution in a particular direction is defined as the full width at half maximum (FWHM) through the center of the system PSF in that direction (Fig. 11b). Lateral and axial resolutions are defined with respect to the collection objective, as is common for LSM.

**Fig. 11.**
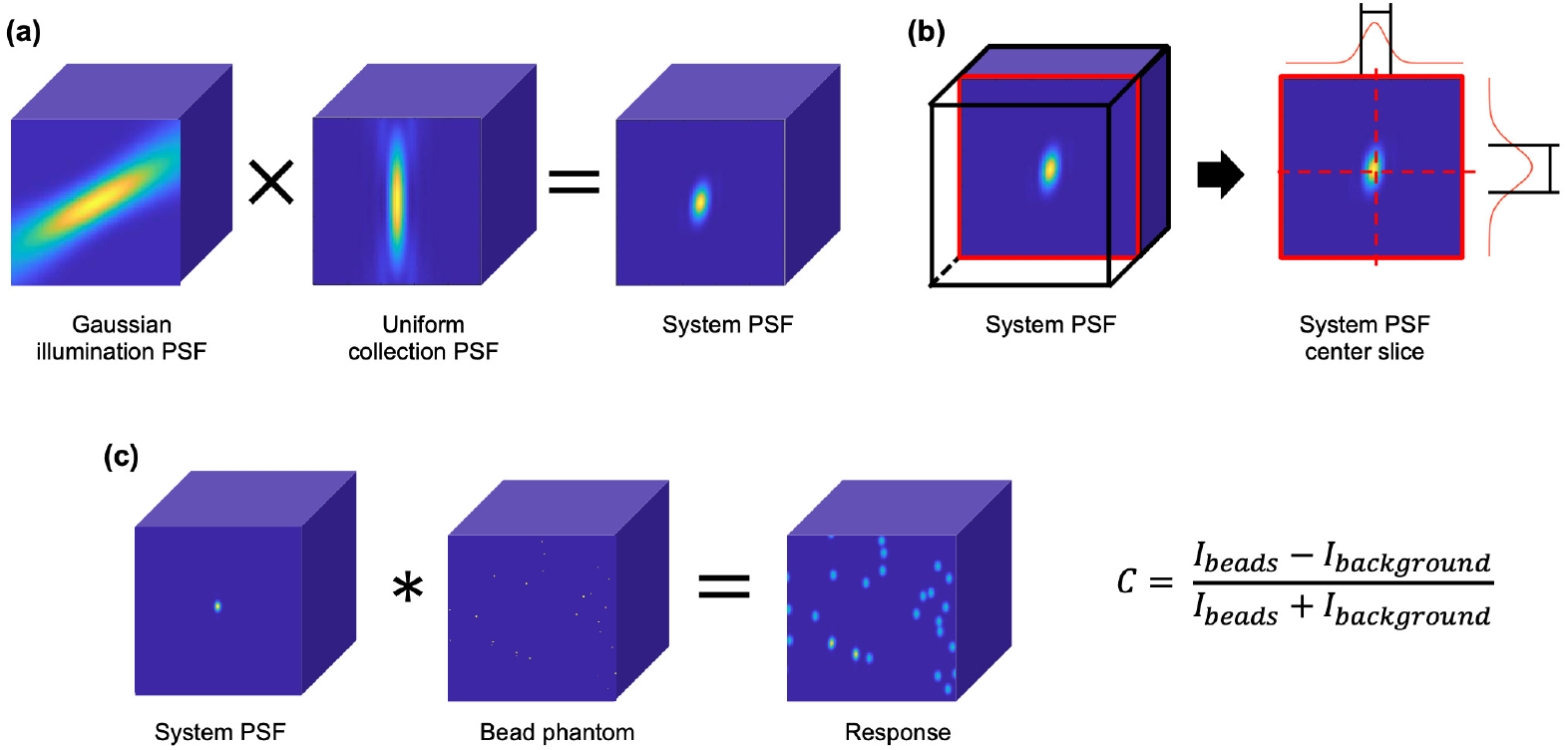
Computational methods for analyzing the resolution and contrast of a given configuration. (a) Three-dimensional point spread functions (PSFs) (assuming Gaussian illumination and uniform collection) are computed separately and multiplied together using the adjoint method to calculate a system PSF. (b) Resolution is computed by finding the full width at half maximum through the center of the system PSF in the lateral and axial directions (relative to the collection objective). (c) To analyze relative differences in image contrast, the system PSF is convolved with a random bead phantom, in which single voxels are set to a value of 1 to simulate diffraction-limited fluorescent beads. Relative contrast levels are then computed based on the value of the response at the locations of the beads (*I_beads_*) relative to the background signal level seen in the images (*I_background_*).

#### Contrast computation

Contrast was computed by simulating the system’s response to a diffraction-limited fluorescent bead phantom. A 3D volume with a voxel pitch of 100 nm was generated in which 0.5% of the voxels were randomly selected as “beads” and set to a value of 1, while all other voxels were set to a value of 0. The normalized system PSF was then convolved with this bead phantom to simulate the system’s response (Fig. 11c). Contrast was computed as:

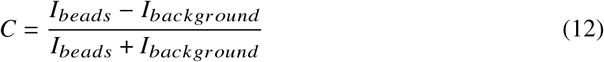

Here, C is contrast, *I_beads_* is the intensity of the response averaged over all the bead locations, and *I_background_* is the average intensity of the response over the bottom 5% of pixels, which adequately represents the background signal for this relatively sparse phantom. Figure 12 shows an example phantom and response with a 0.01% fluorescent bead concentration.

**Fig. 12.**
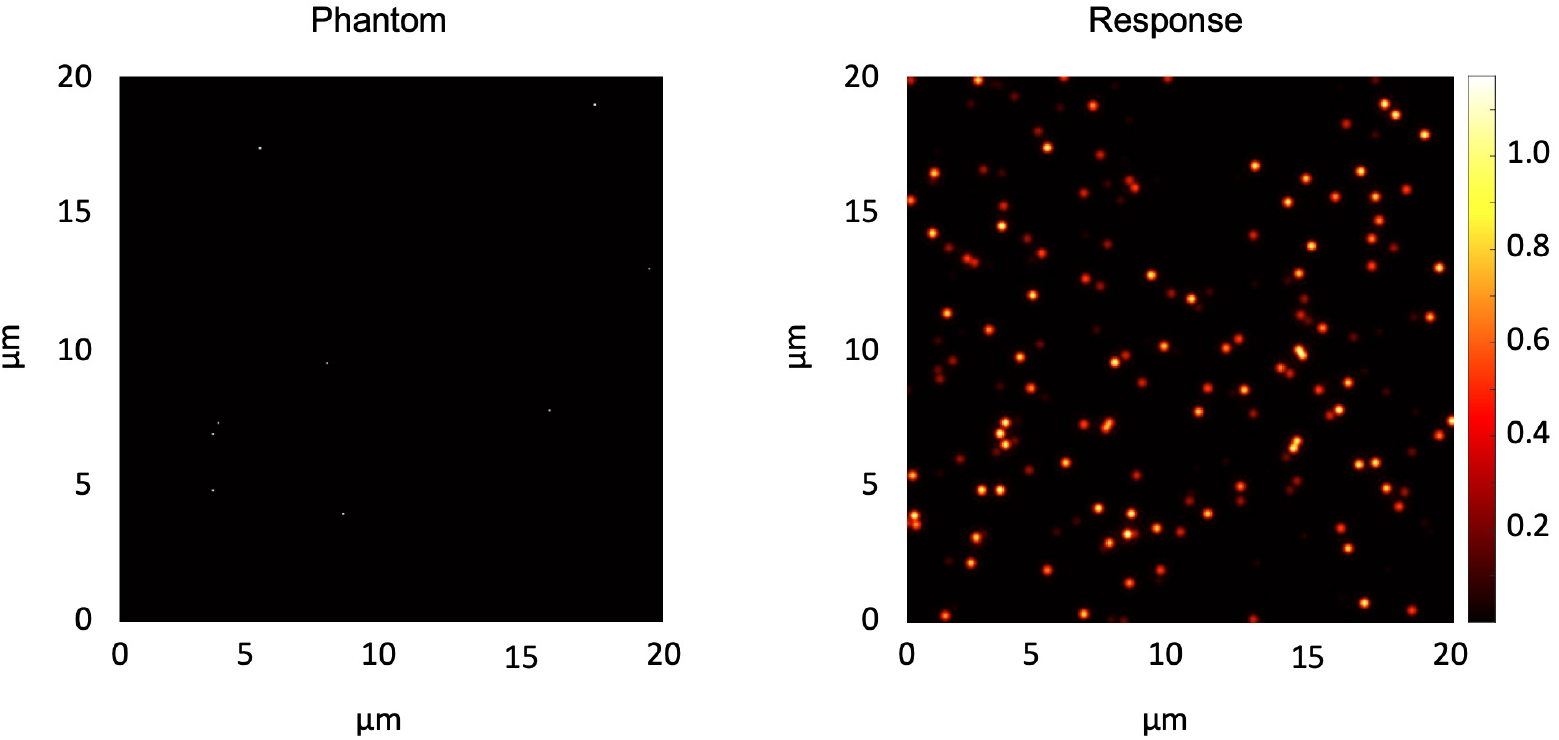
Example diffraction-limited fluorescent bead phantom and response for contrast computation. A 0.01% bead concentration is shown for an orthogonal dual-objective (ODO) system with 0.7 collection NA and 0.2 illumination NA. Note that the figure shows a single slice of a 3D phantom, so many beads visible in the response cannot be seen in the phantom as they are in planes in front of or behind the plane shown.

### Appendix C: Survey of commercial objectives

In order to establish an appropriate range of objective NAs to consider (collection objective NA for dual-objective systems and shared primary objective NA for single-objective systems), we surveyed relevant commercial objectives. As it would not be feasible to consider every existing microscope objective, we limited our search to those designed for, or amenable to, cleared-tissue imaging. Specifically, we considered immersion objectives compatible with refractive indices between 1.38 and 1.56, a range covering most popular clearing protocols [11]. In addition, we only considered objectives with a working distance of at least 1 mm, as shorter working distances would be impractical for volumetric imaging of many cleared-tissue specimens. Finally, we only included objectives with an NA of at least 0.4 (~1 μm or better lateral resolution). Note that some specifications (magnification, NA) change with immersion index for multi-immersion objectives. In these cases, the manufacturer’s nominal specification is used. The objectives identified that meet these criteria are shown in Table 1 and range in NA from 0.4 to 1.0. As such, we used this range for our analysis.

**Table 1.** Survey of commercial objectives for cleared tissue imaging

| Manufacturer | Model | Mag | NA | WD (mm) | Index min | Index max |
| --- | --- | --- | --- | --- | --- | --- |
| LaVision BioTec | 12x NA 0.53 MI Plan - DC33 WD8.5 W | 12x | 0.53 | 8.5 | 1.33 | 1.41 |
| LaVision BioTec | 12x NA 0.53 MI Plan - DC49 WD10.9 AB | 12x | 0.53 | 10.9 | 1.42 | 1.48 |
| LaVision BioTec | 12x NA 0.53 MI Plan - DC57 WD10 O | 12x | 0.53 | 10 | 1.49 | 1.57 |
| Leica | HC FLUOTAR L 25x/1.00 IMM | 25x | 1.0 | 6 | 1.46 | 1.46 |
| Leica | HCX APO L 20x/0.95 IMM | 20x | 0.95 | 1.95 | 1.56 | 1.56 |
| Nikon | CFI PLAN APO 10XC Glyc | 10x | 0.5 | 5.5 | 1.33 | 1.51 |
| Nikon | CFI90 20XC Glyc | 20x | 1.0 | 8.2 | 1.44 | 1.50 |
| Olympus | XL PLN 10X SVM P | 10x | 0.6 | 8 | 1.33 | 1.52 |
| Olympus | XLSLPLN25XGMP | 25x | 1.0 | 8 | 1.41 | 1.52 |
| Olympus | XLSLPLN25XSVM P2 | 25x | 0.95 | 8 | 1.33 | 1.40 |
| Olympus | XLPLN25XSVM P2 | 25x | 1.0 | 4 | 1.33 | 1.40 |
| Special Optics (ASI) | 54-10-12 | 17x | 0.4 | 12 | 1.33 | 1.56 |
| Special Optics (ASI) | 54-12-8 | 24x | 0.7 | 10 | 1.33 | 1.56 |
| Zeiss | Clr Plan-Apochromat 10x/0.5 nd = 1.38 | 10x | 0.5 | 3.7 | 1.38 | 1.38 |
| Zeiss | Clr Plan-Apochromat 20x/1.0 Corr nd = 1.38 | 20x | 1.0 | 5.6 | 1.38 | 1.38 |
| Zeiss | Clr Plan-Neofluar 20x/1.0 Corr nd = 1.45 | 20x | 1.0 | 5.6 | 1.45 | 1.45 |
| Zeiss | Clr Plan-Neofluar 20x/1.0 Corr nd = 1.53 | 20x | 1.0 | 6.4 | 1.53 | 1.53 |

### Appendix D: Contrast of a dual-objective system at various collection NAs

As shown in the Analysis results section, the contrast of a dual-objective system increases with crossing angle until around 60°. Figure 13 shows that this trend holds for other collection NAs (0.4 and 1.0). The plot for a 0.7 collection NA is also reproduced here. Note that there is slight improvement in contrast overall at higher collection NAs. This is because higher collection NAs provide some degree of additional optical sectioning, which makes the PSF in the axial direction narrower and provides a slight boost in contrast.

**Fig. 13.**
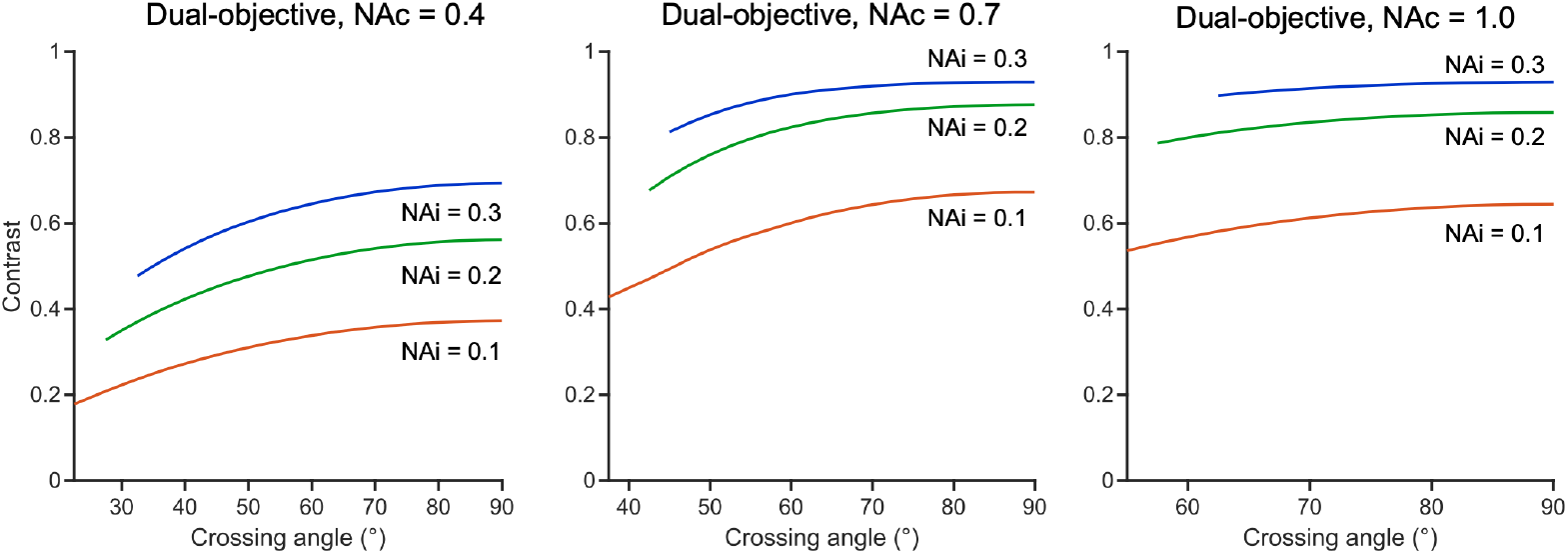
Contrast of a dual-objective system as a function of crossing angle for collection NAs of 0.4, 0.7, and 1.0. In all cases, contrast improves with crossing angle until around 60°, at which point further gains are minimal. Contrast also improves slightly at higher collection NAs, as the collection objective provides some additional optical sectioning at higher NAs.

## Acknowledgments

Any opinions, findings, and conclusions or recommendations expressed in this material are those of the author(s) and do not necessarily reflect the views of the NSF, NIH, or DoD.

